# Evolutionary insights into *Bean common mosaic necrosis virus and Cowpea aphid borne mosaic virus* using global isolates and thirteen new near complete genomes from Kenya

**DOI:** 10.1101/266858

**Authors:** James M Wainaina, Laura Kubatko, Jagger Harvey, Elijah Ateka, Timothy Makori, David Karanja, Laura M. Boykin, Monica A. Kehoe

## Abstract

Plant viral diseases are one of the major limitations in legume production within sub Saharan Africa (SSA), as they account for up to 100 % in production losses within smallholder farms. In this study, field surveys were conducted in the western highlands of Kenya with viral symptomatic leaf samples collected. Subsequently, next-generation sequencing was carried out. The main aim was to gain insights into the selection pressure and evolutionary relationships of *Bean common mosaic necrosis virus* (BCMNV) and *Cowpea aphid-borne mosaic virus* (CABMV), within symptomatic common beans and cowpeas. Eleven near complete genomes of BCMNV and two for CABMV sequences were obtained from SSA. Bayesian phylogenomic analysis and tests for differential selection pressure within sites and across tree branches of the viral genomes was carried out. Three distinct well-supported clades were identified across the whole genome tree, and were in agreement with individual gene trees. Selection pressure analysis within sites and across phylogenetic branches suggested both viruses were evolving independently, but under strong purifying selection, with a slow evolutionary rate. These findings provide valuable insights on the evolution of BCMNV and CABMV genomes and their relationship to other viral genomes globally. These results will contribute greatly to the knowledge gap surrounding the phylogenomic relationship of these viruses, particularly for CABMV, for which there are few genome sequences available, and support the current breeding efforts towards resistance for BCMNV and CABMV.

## 1. Introduction

*Bean common mosaic necrosis virus* (BCMNV) and *Cowpea aphid-borne mosaic virus*(CABMV) belong to the genus *Potyvirus* (Wylie *et al.*, 2017) and are some of the most economically important viruses affecting *Fabaceae* (*Leguminosae*). They have a wide host range and are reported in both cultivated and native vegetation (Worrall et al. 2015; Li et al. 2014). Production losses attributed to BCMNV are up to 100% while for CABMV losses range from 30-60% (Damayanti et al. 2008; Mangeni et al. 2014; Damiri, Abdalla and Amer,2013). Transmission of these viruses are via infected seed stocks they are also transmitted through viruliferous aphids (Worrall *et al.*, 2015).

BCMNV and CABMV are monopartite, single-stranded positive-sense RNA viruses with an average genome size of ~10 Kb and a polyA tail on the 3’-terminal end (El-sawy, Abd and Mohamed, 2014; Fang and Allison, 1995; Mlotshwa et al. 2002). They encode a single polyprotein that is processed into ten proteins in a similar order and function to all members of the family *Potyviridae:* P1 (Symptomatology), HcPro (Aphid transmission), P3 (Plant pathogenicity), 6K1, CI (ATPase/RNA helicase, cell to cell movement), 6K2 (Anchoring the viral replication complex to membranes), Nla-Vpg, Nla-Pro Nlb (Genome replication, RNA-dependent RNA polymerase (RdRp)) and coat protein (CP) (Aphid transmission, virus assembly, cell to cell and systemic movement). There is also a second open reading frame called the PIPO (virulence dependency) (Kehoe et al. 2014; Urcuqui-inchima and Haenni, 2001; Cui et al. 2017; Wei et al. 2010). The small nature of the viral genome and the broad range of environments in which they circulate serve as strong drivers of their genome evolution (Chare, Holmes and Holmes, 2004)

RNA viruses undergo multiple mutations during the replication phase, thus their evolutionary rates (sub/site/year) are considerably higher than cellular genes within their host (Duffy, Shackelton and Holmes, 2008; Holmes, 2009). These nucleotide mutations may result in no change to the amino acids (synonymous changes (dS)), or a change in the amino acid (nonsynonymous changes (dN)). The relative magnitude of these types of changes can lead to positive selection (dN/dS > 1), purifying selection (dN/dS < 1) or neutral selection (dN/dS = 1) within the genome (Nielsen and Yang, 1997; Zhou, Gu and Wilke, 2010). These changes can be quantified to determine the evolution of the virus. Rapid evolutionary rates can lead to new viral strains circulating, while slower evolution rates provide stability in the viral population. Another process that of recombination can also has a profound effect on the molecular evolution of viruses. Particularly in the potyviridae family it is a relatively common occurrence within species, and sometimes even between species (Chare and Holmes 2006; Revers et al 1996; Karasev and Gray 2013; Kehoe et al. 2014)

There have been no studies in Kenya on the molecular evolution of BCMNV and CABMV, particularly within the western highlands of Kenya. The majority of studies in this region are based on serological assays and biological characterisations (Mangeni *et al.*, 2014). Beans (*Phaseolus vulgaris*) and cowpea (*Vigna unguiculata*) play an important role as part of a mixed cropping system for smallholder farmers in Africa, and their role in providing nutrition is crucial. Firstly, they are a source of proteins and micronutrients (iron) especially for women and children in low-income households. Secondly, over the last decade there has been intensive advocacy in the use of legumes as intercrops with cereals such as; maize, sorghum, millet and cassava, which are prioritized as important crops both in Kenya and in the larger sub-Saharan Africa (SSA) region. This has resulted in enhanced interregional seed trade between smallholder farmers and within neighboring countries, with minimal phytosanitary regulation (Katungi *et al.*, 2009). In addition, legumes are currently utilised for integrated pest management of whitefly and whitefly-transmitted viruses (Fondong *et al.*, 2000; Lapidot *et al*, 2014; Uzokwe *et al*, 2016).. A major limitation of this approach has been the limited number of attempts to characterise viruses within these intercropping systems, despite the losses they are capable of causing.

On the basis of this background and in order to fill the existing research gaps on these viruses, we sought to answer the following questions:

1. What are the evolutionary relationships of *bean common mosaic necrosis virus* and *cowpea aphid borne mosaic virus*?
2. What selective pressures are these viruses under and how do they govern the evolution of these viruses?

To answer these questions we used a viral metagenomic approach using next generation sequencing (NGS) on symptomatic beans and cowpeas. NGS applications in plant virology began in 2009 (Blawid, Silva and Nagata, 2017) and have been increasingly applied in the *de novo* discovery of RNA and DNA viruses as well as viroids due to its rise as a rapid and relatively inexpensive mode of viral detection (Hadidi et al. 2016; Blawid, Silva and Nagata, 2017). Previous reports of using NGS for novel viral discovery and subsequent evolutionary analysis are well-documented (Kehoe et al. 2014; Alicai et al. 2016; Ndunguru et al. 2015; Wamonje et al. 2017; Maina et al. 2017; Kraberger et al. 2017). In this study, we use this approach to obtain near complete genomes of BCMNV and CABMV, and gain further insights into the evolutionary relationships of BCMNV and CABMV.

## 2 Materials and methods

### 2.1. Field collection

Field surveys were carried out within the highlands of western Kenya over two cropping seasons (2015 and 2016) during the long rains. Sampling was conducted in heterogeneous cropping systems predominately comprising beans (*Phaseolus vulgaris L*) (from Busia, Bungoma, Kakamega and Vihiga counties) and cowpea *(Vigna unguiculata (L.)*Walp.) (from Busia). Leaf samples collected from symptomatic plants were stored using two methods: silica gel and the paper press method (Abu Almakarem *et al.*, 2012). Samples were then transported to the Bioscience eastern central Africa (BecA-ILRI) laboratories in Nairobi, Kenya for processing.

### 2.2. Nucleic acid extraction and PCR screening viruses

From each individual leaf, RNA was extracted using the Zymo RNA miniprep kit (Zymo, USA) according to the manufacturer’s specifications. Extractions were then lyophilised and shipped to the University of Western Australia for further processing. Screening for positive *Potyvirus* samples was done using the universal *Potyvirus* primer LegPotyF 5’-GCWKCHATGATY GARGCHT GGG-3’ and LegPotyR 5’-AYYTGYTYMTCHCCATCCATC-3’ (Webster, 2008).

### 2.3. cDNA library preparation and RNAseq sequencing

A total of 28 individual cDNA libraries of each of the samples (n=24 beans, n= 4 cowpea) were prepared using Illumina Truseq stranded total RNA sample preparation kit with Plant Ribozero as described by the manufacturer (Illumina). All libraries containing the correct insert size fragment and quantity were sent to Macrogen Korea for subsequent sequencing. Libraries were normalised based on the concentration and then pooled prior to sequencing. Pair end sequencing (2 × 150 nt) was done on the rapid run mode using a single flow cell on the Illumina Hiseq 2500.

### 2.4. Assembly and mapping of RNAseq reads

Raw reads were trimmed and assembled using CLC Genomics workbench (CLCGW ver 7.0.5) (Qiagen). Trimmed reads were assembled using the following parameters: quality scores limit set to 0.01, maximum number of ambiguities was set to two and read lengths less than 100 nt were discarded. Contigs were assembled using the *de novo* assembly function on CLCGW with default automatic word size, and automatic bubble size parameters. Minimum contig length was set to 500, mismatch cost two, insertion cost three, deletion cost three, length fraction 0.5 and similarity fraction 0.9. All the contigs were subjected to Blastn and Blastx (NCBI) on the Magnus Supercomputer at Pawsey. Contigs that matched plant viruses were identified and exported to Geneious 8.1.5 (Biomatters). Reference-based mapping was then carried out using complete genomes retrieved from GenBank reference (KX302007 for BCMNV, while for CABMV the closet complete match was MF179118 (this study)). Mapping parameters were set as follows: minimum overlap 10%, minimum overlap identity 80%, allow gaps 10% and fine tuning iteration up to 10 times. The consensus contig from the mapping was aligned using MAFFT (Katoh and Standley 2013) to the *de novo* contig of interest. The resulting alignments were manually inspected for ambiguities, which were corrected with reference to the original assembly or mapping. The open reading frame and annotation of the final sequences was done in Geneious 8.1.5 (Biomatters). Sequences were referred to as nearly complete if the entire coding region was present, and complete if the entire genome including untranslated regions were present.

### 2.5. Detection of recombination break points

Assessment of the recombination break points of the nearly complete genomes of BCMNV (n =11) and CABMV (n=2) from this study, and those retrieved from GenBank (BCMNV n= 9 and CABMNV n= 6) was carried out using the seven programs within the RDP4 software (Martin *et al.*, 2015). The programs used were: RDP (Martin *et al*, 2005), GENECONV (Padidam, Sawyer and Fauquet, 1999), Bootscan (Martin *et al.*, 2005) MaxChi (Smith, 1992) Chimaera (Posada and Crandall 2001), 3Seq (Boni, Posada and Feldman, 2007) and SiScan (Gibbs, Armstrong and Gibbs, 2000). A recombination event was detected only if found by at least four of the seven programs and further supported by a Bonferroni correction with a *P* value cut-off of 0.05.

### 2.6. Bayesian phylogenetic analysis of BCMNV and CABMV

Bayesian inference was used to establish the phylogenetic relationships within BCMNV and CABMV from this study with *Bean common mosaic virus* (BCMV) as the outgroup. The analysis was carried out on both the near complete genomes and separately on the ten individual genes. The most suitable evolutionary models were determined by jModelTest (Darriba *et al*, 2012) and bModelTest (Bouckaert and Drummond 2017). Bayesian analysis of the nearly complete genomes was carried out using Exabayes 1.4.1 (Aberer, Kobert and Stamatakis, 2014) while individual genes were analysed using MrBayes 3.2.2 (Huelsenbeck, 2001). Exabayes was selected for the genome analysis since it can assign independent evolutionary models to each of ten individual genes within a single run, while MrBayes utilizes a single evolutionary model for each gene in a given run. MrBayes was run for 50 million generations on four chains, with trees sampled every 1000 generations using GTR+I+G as the evolutionary model. In each of the runs, the first 25% (2,500) of the sampled trees were discarded as burn-in. In the ExaBayes run, each gene segment was assigned an independent evolutionary model. ExaBayes was run was for 50 million generations on four chains. In each run, the first 25% of the sampled trees were discarded as burn-in. Convergence and mixing of the chains was evaluated using Tracer v1.6 (http://tree.bio.ed.ac.uk/software/tracer/)

### 2.7. Analysis of extent of selection pressure across sites with whole genome

Selective pressure was evaluated by dN/dS, across sites of the coding region using SNAP (Aberer, Kobert and Stamatakis, 2014). Sites under positive selection were identified using SLAC in Datamonkey (http://www.datamonkey.org/).

### 2.8. Analysis of selection pressure

Selection pressure is analysed using the nonsynonymous to synonymous substitution rate ratio, or ω (ω = d_N_/d_S_). For BCMNV and CABMV, this was done using a maximum likelihood codon-substitution model in the CODEML program in PAML 4 (Yang, 2007). A representative subset (n = 14), (n = 10 from this study and n = 4 representative from GenBank) from each of the three clades of the phylogenetic tree obtained above was used in the analysis. Two models were assessed by a likelihood ration test (LRT): i) M2a vs M1a (neutral vs positive selection) ii) M3 vs M0 (variable ratio vs one single ratio).

### 2.9. Estimation of selection pressure between clades of the gene trees

Differences in the selection pressure between the two main clades I and II (Fig. 2A-2J) were assessed by testing the null hypothesis of equal selection pressure (Model = 0 Nsite = 0) against the alternative hypothesis that each clade has a distinct to (Model = 1 Nsite = 0, different selection pressure) across all ten gene trees. The test for significance was based on the likelihood ratio test (LRT) with significance level at α = 0.05.

## 3. Results and discussion

### 3.1. Sample screening, next generation sequencing and data assembly

This study provides insights into the phylogenetic relationships of BCMNV and CABMV and selective pressures that govern their evolution. We present the near complete genomes of eleven BCMNV and two whole genomes of CABMV from Kenya. These viruses were under strong purifying selection with BCMNV and CABMV evolving independently and at a slow rate. These genomes are the first from SSA and are an invaluable genomic resource to not only help understand the molecular evolution of these viruses but also assist in the development of molecular diagnostic tools.

Legume samples that tested positive using universal primers for *Potyvirus* were selected for library preparation and subsequent next generation sequencing. Samples that were negative after PCR amplification were excluded from library preparation. These negative samples were probably due to nutritional deficiency rather than viral infections. RNAseq of total plant RNA resulted in raw reads (after trimming) that ranged between 12,667,976 and 15,638,762 reads. *De novo* assembly produced a number of contigs ranging between 895-19,029 nucleotides (nt) (Table S2). Plant virus contigs were identified after BLASTn searches with lengths of between 1,048 − 9,763 nt, and average coverage depth of between 71 − 8,924 times. Genomes with the complete open reading frame and including the complete untranslated regions were considered to be full genomes. However, genome sequences that lacked parts of the 5’ and 3’ UTR regions were referred to as near complete genomes. The final sequence was obtained from the consensus of *de novo* assembly and the mapped consensus of reads and ranged from 8,241 to 9,837 nt in length (Table S1). BCMNV and CABMV sequences obtained from this study are summarised in Table 1, while sequences retrieved from GenBank and associated metadata are provided in Table S1. In total, there were 11 BCMNV and two CABMV genomes from this study (Table 1). All viral sequences generated from this study were deposited in GenBank with the accession numbers MF179108 − MF179120.

**Table 1.** Near complete genomes of *Bean common mosaic necrosis virus* (BCMNV) and the *Cowpea aphid-borne mosaic virus* (CABMV) collected across two seasons 2015/2016 from the western highlands of Kenya

| Sample ID | GenBank | Virus Identified | Plant Host | Geographical location | Collection Year |
| --- | --- | --- | --- | --- | --- |
| SRF 08 | MF179117 | BCMNV | Bean | Kakamega | 2015 |
| SRF 33 | MF179112 | BCMNV | Bean | Kakamega | 2015 |
| SRF 50 | MF179115 | BCMNV | Bean | Kakamega | 2016 |
| SRF61 | MF179109 | BCMNV | Bean | Busia | 2016 |
| SRF 74 | MF179120 | CABMV | Cowpea | Bungoma | 2016 |
| SRF 75 | MF179111 | BCMNV | Bean | Bungoma | 2016 |
| SRF 77 | MF179118 | CABMV | Cowpea | Bungoma | 2016 |
| SRF 97 | MF179113 | BCMNV | Bean | Kakamega | 2016 |
| SRF 99 | MF179110 | BCMNV | Bean | Kakamega | 2016 |
| SRF 114 | MF179116 | BCMNV | Bean | Vihiga | 2016 |
| SRF 119 | MF179114 | BCMNV | Bean | Vihiga | 2016 |
| SRF 122 | MF179108 | BCMNV | Bean | Vihiga | 2016 |

### 3.2. Analysis of recombination analysis of BCMNV and CABMV

Two BCMNV isolates (Table 2) and three CABMV isolates (Table 3) from Kenya were identified as recombinant. None of the CABMV recombinants sequences were from this study. The locations of recombination were within the P1/HcPro/P3 in (BCMNV) and Nla Pro/Nlb/CP regions in (CABMV) (Table 2 and 3). These sequences were excluded from subsequent analysis. One of the drivers of viral evolution is recombination (Roossinck, 2003; Valli, López-Moya and García, 2007). Previous reports have indicated that BCMV-BCMNV recombination occurs resulting in new stable and virulent BCMNV strains (Larsen *et al.*,2005). Similar findings have been reported in other RNA viruses such as *Turnip mosaic virus* (Nguyen *et al.*, 2013), *Papaya ring spot virus* (Maina *et al.*, 2017) and *Cassava brown streak virus* (Ndunguru *et al.*, 2015). Since, recombinant sequences can distort the true relationships when studying the phylogenetic relationships between sequences, we excluded all recombinants from subsequent downstream analysis (Varsani et al. 2008; Penny et al. 2008; Posada and Crandall, 2001). In addition, during the library preparation individual samples were used for library preparation and each was dual-indexed, ensuring improved integrity of the viral genomes recovered and used for phylogenetic analysis.

**Table 2.** Recombination signals across BCMNV using RDP4. Table entries represent the recombinant sequences and the position of recombination within the complete genome. A recombinant was considered as true recombinant if more than four detection programs supported at a significance level of 0.05.

| Recombination Events | Recombinant Sequence | Detected Breakpoint | Parental Sequence (Major) | Parental Sequence (Minor) | Detected in RDP4 | Avr P-Val |
| --- | --- | --- | --- | --- | --- | --- |
| 1 | SRF08_ MF179117 | 5515-8886 | Kenya_Beans_SRF50 | USA_Beans_HQ229994 | R, M, C, S , 3 | 3.55 X 10<sup>-4</sup> |
| 2 | SRF114_ MF179116 | 1341-1619 | Kenya_Beans_SRF50 | USA_Beans_HQ229993 | R, G, B, C M 3 | 0.0415 |

**Key:** Recombinant programs in RDP4 that detected recombinant events across the whole genome of BCMNV
3=3seq B=Bootscan C=Chimera G=Gencov R=RDP M=Maxchi S=Siscan

**Table 3.** Recombination signals across CABMV using RDP4 (Martin *et al.*, 2015). Table entries represent the recombinant sequences and the position of recombination within the complete genome. A recombinant was considered as true recombinant if more than four detection programs supported at a significance level of 0.05.

| Recombination event | Recombinant Sequence | Detected Breakpoint | Parental Sequence (Major) | Parental Sequence( Minor) | Detected in RDP4 | Aver P Value |
| --- | --- | --- | --- | --- | --- | --- |
| 1 | Brazil_ HQ880243 | 2332-2396 | Zimbabwe_NC004013.1 | Brazil_Cowpea_HQ880242 | R,G, B,M,C,S,**3** | 2.442 X 10<sup>-14</sup> |
| 2 | India_ KM597165 | 917-1220 | Brazil_HQ880242 | Brazil_Cowpea_HQ880243 | R, G, B, M, C, **S**, 3 | 1.800 X 10<sup>-14</sup> |
| 3 | Brazil_ HQ880242 | 1385-1409 | India_KM597165 | Brazil_Cowpea_HQ880243 | R, G, B, M, C,**S**,3 | 1.571 X10<sup>-07</sup> |
| 6 | Brazil_ HQ880243 | 8643-8809 | India_ KM597165 | Uganda_Cowpea_KT726938 | R, G, B, **M**, C, 3 | 1.328 X 10<sup>-04</sup> |
| 7 | Brazil_ HQ880242<br>India_ KM597165 | 94-275 | Zimbabwe_ NC004013.1 | Uganda_Cowpea_KT726938 | R, **G**, B, M, C,3 | 4.750 X 10<sup>-4</sup> |
| 10 | Brazil_ HQ880243 | 6605-8134 | Zimbabwe_ NC004013.1 | Brazil_Cowpea_HQ880242 | R, B, M, **C**,S | 2.595 X 10<sup>-2</sup> |
| 18 | Brazil_ HQ880243<br>Brazil_ HQ880242<br>India_ KM597165 | 6916-8065 | Uganda_ KT726938 | Zimbabwe_Cowpea_NC004013.1 | R, **B**, M, C, S | 1.664 X 10<sup>-2</sup> |

**Key:** Recombinant programs in RDP4 that detected recombinant events across the whole genome of CABMV **3**=3seq **B**=Bootscan **C**=Chimera **G**=Gencov **R**=RDP **M**=Maxchi **S**=Siscan

### 3.3 Bayesian evolutionary relationship of BCMNV and CABMV

Phylogenetic relationships within BCMNV and CABMV were based on the near complete genome tree (Fig. 1) and with reference to the ten individual gene trees (Fig. 2). Both the whole genome tree and the gene trees resulted in identical topologies. Two well-supported clades I and II within BCMNV and only one clade within CABMV were identified, which we have called III for the purposes of this research (Fig. 1 and Fig. 2). The individual gene trees resulted in two main clades within the BCMNV sequence, and CABMV sequences formed a single monophyletic group (Fig. 1). Across the two BCMNV clades (I and II) the percentage nucleotide identities were 96.1 − 98.9 % as we would be expected for member of the same potyvirus species. Likewise, the CABMV clade (III) showed 79.3% nucleotide identity, as expected for within species comparison. However, CABMV showed species diversity when compared to the BCMNV species (Table 4).

**Fig. 1.**
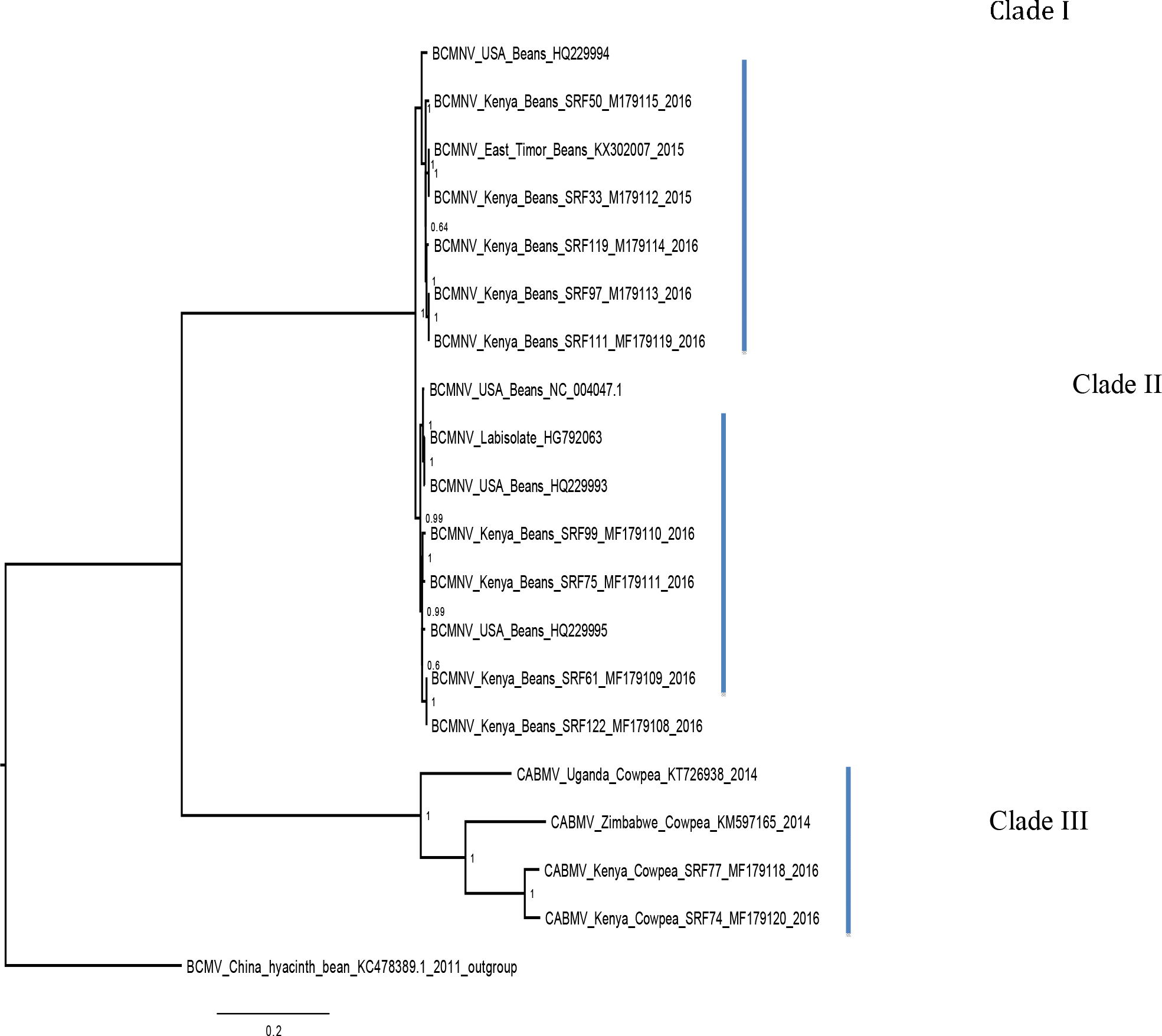
Consensus of trees sampled in a Bayesian analysis of the whole genome of *Bean common mosaic necrosis virus* (BCMNV) and *Cowpeas aphid borne mosaic virus* (CABMV) with *Bean common mosaic virus* (BCMV) used as an outgroup using ExaBayes 1.4.1

**Fig. 2.**
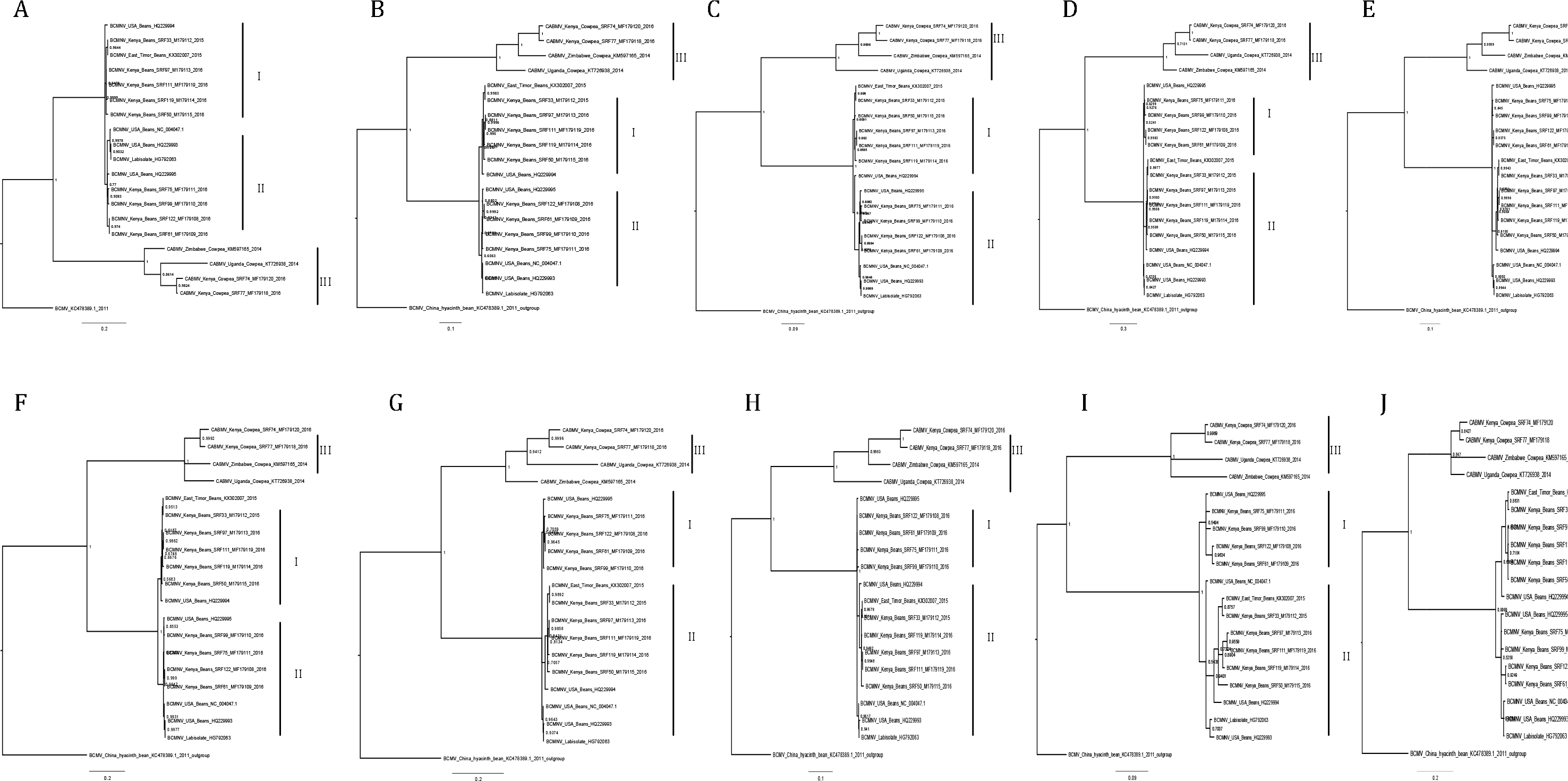
Consensus of trees sampled in a Bayesian analysis gene tree of *Bean common mosaic necrosis virus* (BCMNV) and *Cowpeas aphid borne mosaic virus* (CABMV) with *Bean common mosaic virus* (BCMV) used as an outgroup using MrBayes 3.2.2. **(A)**, coat protein (CP) **(B)** CI **(C)** Nlb **(D)** PI **(E)** Hc-Pro **(F)** P3 **(G)** Nla-Pro **(H)** Nla-Vpg **(I)** 6K2 **(J)** 6K1

**Table 4.**
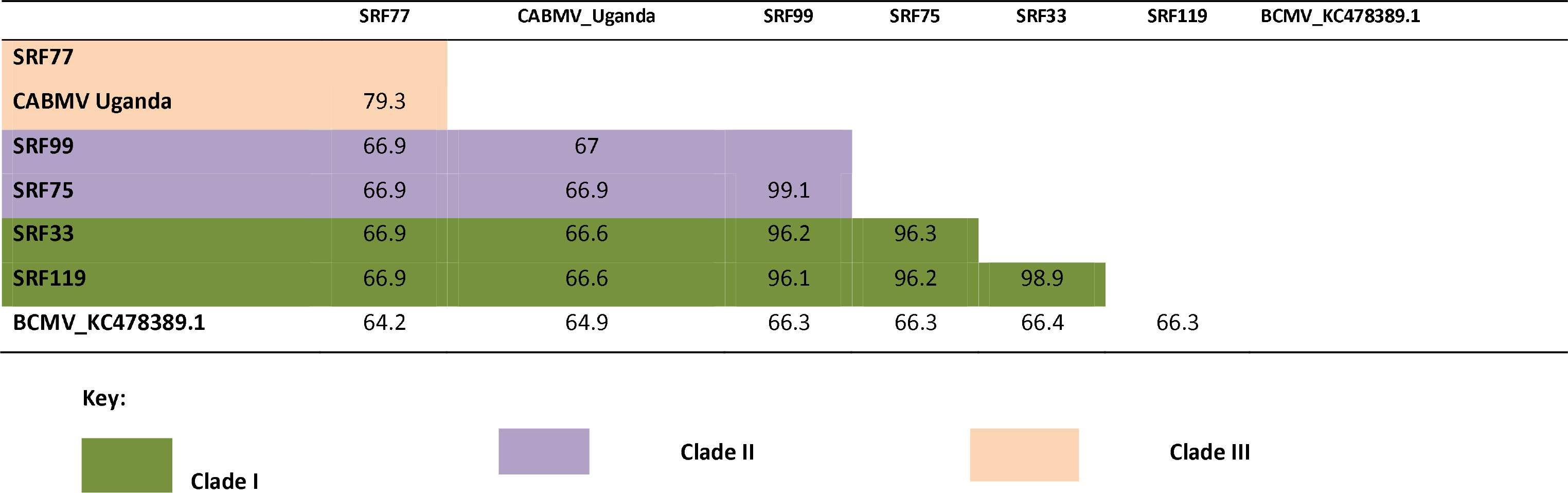
Pairwise sequence comparison of the nearly complete genomes of between two representative sequences across the three clades with BCMV_KC478389.1 as the outgroup using Geneious 8.1.8. Intraclade similarity was over 79 % across all the three clades.

|  | SRF77 | CABMV_Uganda | SRF99 | SRF75 | SRF33 | SRF119 | BCMV_KC478389.1 |
| --- | --- | --- | --- | --- | --- | --- | --- |
| SRF77 |  |  |  |  |  |  |  |
| CABMV Uganda | 79.3 |  |  |  |  |  |  |
| SRF99 | 66.9 | 67 |  |  |  |  |  |
| SRF75 | 66.9 | 66.9 | 99.1 |  |  |  |  |
| SRF33 | 66.9 | 66.6 | 96.2 | 96.3 |  |  |  |
| SRF119 | 66.9 | 66.6 | 96.1 | 96.2 | 98.9 |  |  |
| BCMV_KC478389.1 | 64.2 | 64.9 | 66.3 | 66.3 | 66.4 | 66.3 |  |

Clade I
Clade II
Clade III

We suspect that the drivers of BCMNV and CABMV diversity are similar to other members of the *Potyviridae*. Within *Potyviridae* the evolutionary divergence is thought to occur through the star burst phenomena, that occurs during the introduction of new viruses onto native lands (Gibbs et al. 2008). Native ecosystems act as a catalyst for genetic divergence of these viruses resulting in new viral strains and/or quasispecies. Similarity in tree topologies between the genome tree and the coat protein gene tree indicate the coat protein is a reliable phylogenetic marker (Fig. 1 and Fig. 2a) and the topologies recovered in our analyses are similar to previously reported typologies (Zheng et al. 2002). However, our results differ from previous studies within other members of *Potyviridae* (CBSV, UCBSV), where the whole genome tree, and their coat protein gene tree did not concur with the species trees (Alicai et al. 2016). This highlights the diversity of the members of the *Potyviridae* and the importance of performing rigorous phylogenetic, recombination and evolutionary analysis for each individual species.

### 3.4. Estimation of selection pressure across sites and branches of the whole genome

Episodic positive selection pressure along the branches of the BCMNV, BCMV and CABMV phylogenetic tree (Fig. 2) was estimated using the branch models in CODEML within PAML 4 using two models. Model 2 supported independent episodic changes along the branches of BCMNV, CABMV and BCMV was accepted based on the log likelihood ratios of _p < 0.05_ (Table 5). Thus rejecting the null test model (M_0_) of equal episodic changes within BCMNV, CABMV and BCMV (Table 5). In addition, the selective pressure within the genes of both BCMNV and CABMV supported high purifying selection ω < 1 (Fig. 3) across all genes. However, the site selective pressure as determined by analysis using SNAP was not uniform across the genes (Fig 3). Selective pressure within BCMNV genes was highest in Nlb, Nla-Vpg and lowest in PI and P3 (Fig. 3A). While in CABMV genes, the highest site selective pressure site were 6K1 and Nla-Pro, while the least site pressure was in P1 and P3 (Fig. 3B). The two clades within BCMNV gene trees were under equal selective pressure except within the CI gene (Table 6). These selective pressures have resulted in high purifying selection across the genes and could be responsible in maintaining the function of theses genes.

**Fig. 3.**
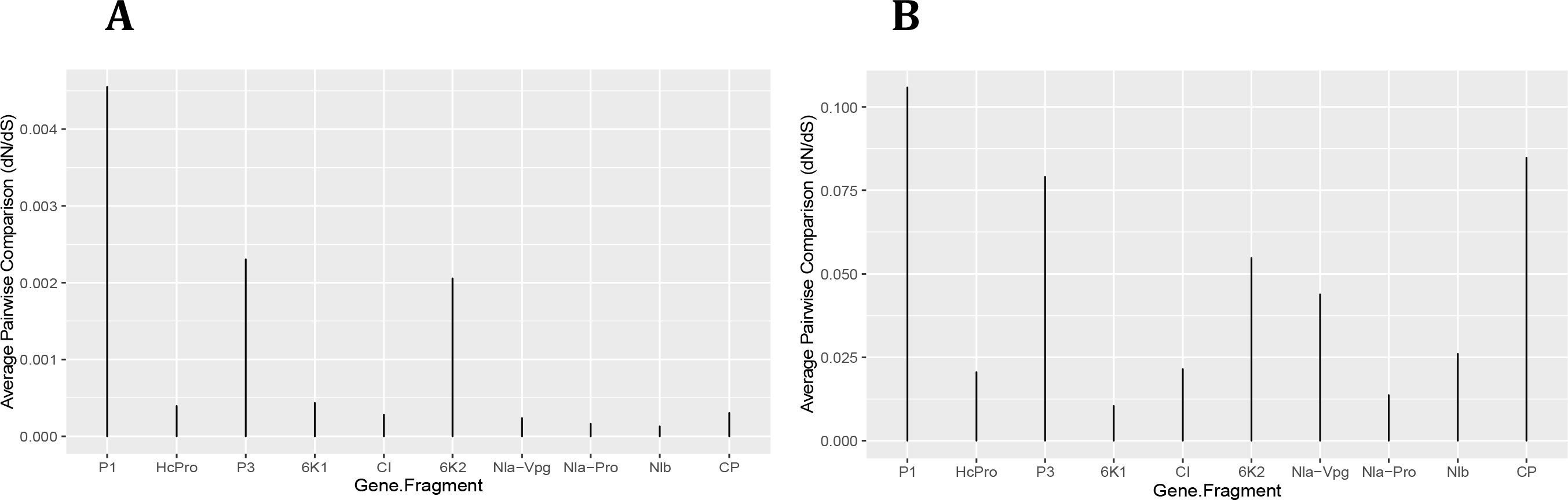
Selection pressure within sites across the viral gene fragments was determined by assessing the average synonymous and non-synonymous (d_N_/d_S_) a the coding region using SNAP that were plotted against each gene viral gene segment subsequently plotted using ggplot in R ver. 0.99. All sites are under s purifying selection with d_N_/d_S_ values < 1. (A) Comparison of the average synonymous (dS) and non-synonymous (dN) sites across the coding region of each segment in BCMNV near complete genome using SNAP. (B) Comparison of the average synonymous (dS) and non-synonymous (dN) across the coding reg each gene segment in CABMV near complete genome using SNAP.

**Table 5.**
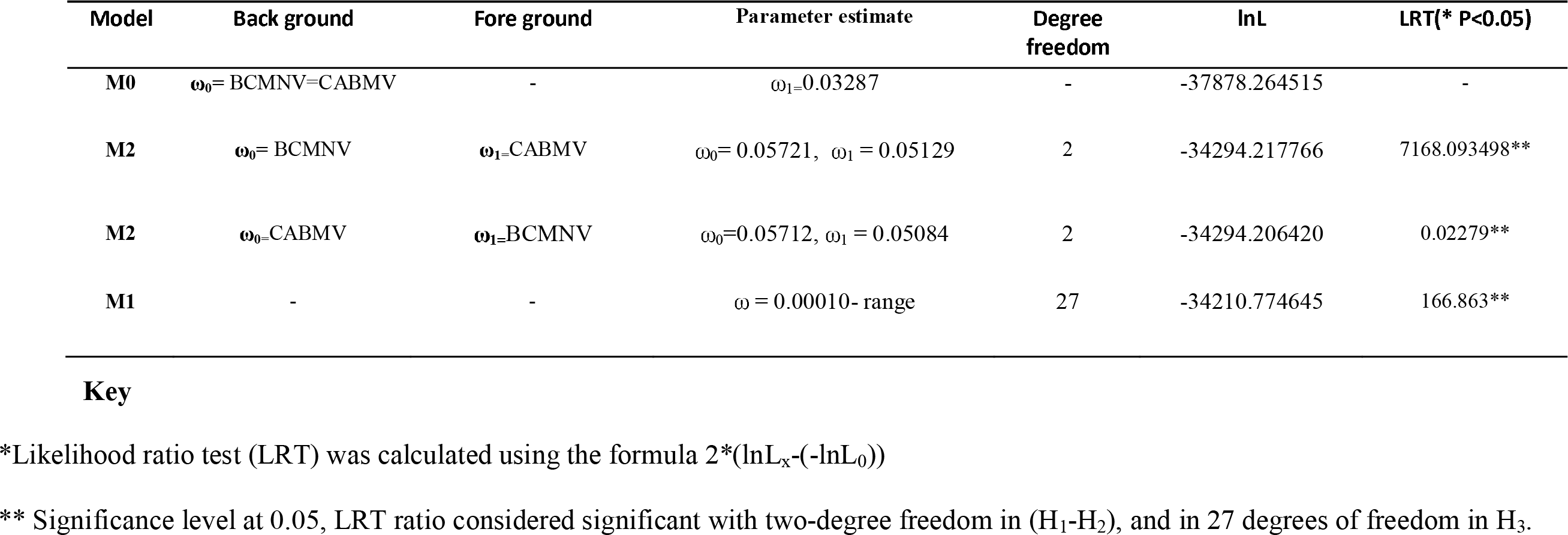
The dn/ds (ω) values, log-likelihood (lnL) values, likelihood ratio test (LRT) statistics and positively selected branches under different models of codon substitution were used to investigate selection pressures on the 8883 nucleotide of BCMNV-BCMV and CABMV analysed in this study

| Model | Back ground | Fore ground | Parameter estimate | Degree freedom | lnL | LRT(* P<0.05) |
| --- | --- | --- | --- | --- | --- | --- |
| M0 | $\omega_0 = \text{BCMNV}=\text{CABMV}$ | - | $\omega_1=0.03287$ | - | -37878.264515 | - |
| M2 | $\omega_0 = \text{BCMNV}$ | $\omega_1=\text{CABMV}$ | $\omega_0 = 0.05721, \omega_1 = 0.05129$ | 2 | -34294.217766 | 7168.093498** |
| M2 | $\omega_0=\text{CABMV}$ | $\omega_1=\text{BCMNV}$ | $\omega_0=0.05712, \omega_1 = 0.05084$ | 2 | -34294.206420 | 0.02279** |
| M1 | - | - | $\omega = 0.00010\text{- range}$ | 27 | -34210.774645 | 166.863** |

\*Likelihood ratio test (LRT) was calculated using the formula $2*(\ln L_x - (-\ln L_0))$
\*\* Significance level at 0.05, LRT ratio considered significant with two-degree freedom in ( $H_1-H_2$ ), and in 27 degrees of freedom in $H_3$ .

**Table 6.** Comparison of rates of evolution within the two clades within BCMNV across the gene fragments. The null hypothesis is that there are equal rates while the alternative specifies different ω values between the two clades. Tests are carried out at a significance level of 0.05.

| Gene | Equal rates (null hypothesis) |  | Differing rates (alternative hypothesis) |  |  | LRT statistic | P value |
| --- | --- | --- | --- | --- | --- | --- | --- |
| | lnL | $\omega$ dN/dS) | lnL | $\omega 0$ | $\omega 1$ | | |
| P1 | -2214.505595 | 0.37095 | -2214.452326 | 0.37610 | 0.30129 | 0.106538 | P > 0.05 |
| HcPro | -2795.992899 | 0.09728 | -2794.811926 | 0.08884 | 0.60647 | 2.361956 | P > 0.05 |
| P3 | -2055.983286 | 0.28547 | -2054.343977 | 0.27297 | 999.00000*** | 3.27862 | P > 0.05 |
| 6K1 | -323.414987 | 0.05870 | -323.205094 | 0.06221 | 0.00010*** | 0.419786 | P > 0.05 |
| CI | -3606.604936 | 0.08320 | -3602.434146 | 0.06552 | 0.32492 | 8.341508 | P < 0.05 |
| 6K2 | -341.075396 | 0.24049 | -341.075137 | 0.24140 | 0.23425 | 0.00052 | P > 0.05 |
| Nla-Vpg | -979.236913 | 0.09398 | -978.201866 | 0.11081 | 0.00010*** | 2.07009 | P > 0.05 |
| Nla-Pro | -1392.256063 | 0.08420 | -1391.975219 | 0.08673 | 0.00010*** | 0.56168 | P > 0.05 |
| Nlb | -2997.347051 | 0.06322 | -2997.340969 | 0.06285 | 0.07124 | 0.012164 | P > 0.05 |
| CP | -735.455148 | 0.09977 | -735.394608 | 0.10664 | 0.07166 | 0.12108 | P > 0.05 |

\*\*\* $\omega$ estimate to be on the boundary of allowable values in PAML

The Nlb, Nla-Vpg in BCMNV and 6K1 and Nla-Pro genes are under high selective pressure for both BCMNV and CABMV are associated with viral genome replication. The continuous survival of viruses is dependent on successful replication, which could be a key driver in ensuring that these genes undergo high purifying selection pressure in spite of recombination and mutations that may occur as observed (Table 2 and 3) to ensure they maintain these critical function. These findings are similar to other vector transmitted RNA viruses such as the *Rice stripe virus* (RSV) (Wei et al. 2008; He et al. 2017). A probable explanation of the high purifying selection is due to the trade-off phenomena (Holmes, 2009; Chare and Holmes, 2004), experienced in vector-transmitted viruses. Vector-transmitted RNA viruses have a wide host range and multiple insect vectors, in this case over 200 aphid species. To increase their chances of multiple vectors transmitting them effectively into several plant hosts, they undergo strong purifying selection resulting in slower rates of evolution. This is contrary to non-vector transmitted RNA viruses that show high selection pressure indicative of positive selection (Wood et al. 2009; Troupin et al. 2016). This further validates that the evolution of vector-transmitted viruses is constrained by their vectors rather than their plant host (Holmes, 2009; Chare and Holmes, 2004). A slower rate in the evolution of BCMNV and CABMV could be beneficial in the breeding efforts, since it would allow for the new BCMNV resistant varieties under development to persist for longer than if the viruses were evolving at faster rates. The importance of continuous monitoring of viral evolution ensures that breeding efforts against viruses remain relevant. This is especially important considering the numerous cases of resistance breakdown associated with continuous virus evolution. There is a direct correlation between the evolution of viruses and the durability of the resistance (García-Arenal and McDonald 2003).

## 4. Conclusions and future perspectives

In this study, we identified two main clades within BCMNV and a single clade within CABMV based on phylogenomic analysis using the whole genome and ten gene trees. The overall evolution rates within BCMNV and CABMV revealed the viruses were under strong purifying selection and thus evolving slowly. These findings provide robust genomic and evolutionary data to complement current bean research underway in Africa at the BecA-ILRI hub complemented by efforts at Cambridge University. In addition, this study highlights the need to establish robust biosecurity and phytosanitary measures within developing countries in order to control the spread of these viruses. That they appear to be relatively stable in an evolutionary sense at this point in time bodes well for plant disease management strategy development. We strongly advocate for the use of cultural control strategies as the first line of defence against these viruses within smallholder farming communities.

## Acknowledgments

J.M.W is supported by an Australian Award scholarship by the Department of Foreign Affairs and Trade (DFAT), this work forms part of his PhD research. Initial laboratory work was performed at the Biosciences eastern and central Africa (BecA-ILRI Hub) Nairobi,Kenya. Fieldwork activities were coordinates through Cassava Diagnostic project Kenyan node. Pawsey Supercomputing Centre provided supercomputer resources for data analysis with funding from the Australian Government and the Government of Western Australia.

## Appendix A. Supplementary Material

Supplementary tables

